# Introducing ExHiBITT – Exploring Host microbiome inTeractions in Twins-, a colon multiomic cohort study

**DOI:** 10.1101/795070

**Authors:** Marina Mora-Ortiz, Hajir Ibraheim, Sherine Hermangild Kottoor, Ruth C. E. Bowyer, Sarah Metrustry, Jeremy Sanderson, Nicholas Powell, Tim D. Spector, Kerrin S. Small, Claire J. Steves

## Abstract

The colon is populated by approximately 10^12^ microorganisms, but the relationships between this microbiome and the host health status are still not completely understood. Participants from the TwinsUK cohort were recruited to study the interactions between the microbiome and host adaptive immunity. In total, 205 monozygotic twins were recruited from the wider TwinsUK cohort. They completed health questionnaires, and provided saliva, blood, colon biopsies from three different locations, caecal fluid, and two faecal-samples.

Here, our objective is to present the cohort characteristics of **ExHiBITT** including i) biomedical phenotypes, ii) environmental factors and ii) colonoscopic findings. A significant proportion of this apparently normal cohort had colonic polyps (28%), which are of interest as potential precursors of colorectal cancer, and as expected, the number of polyps found was significantly correlated with BMI and age. Hitherto undiagnosed diverticulosis was also not infrequently found during colonoscopy (26%) and was associated in changes in Hybrid Th1-17 cells in the colon. Twin proband cooccurrence rate for diverticulosis (82%), was much higher than for polyps (42%). Familial factors affecting morphology or tolerance may contribute to the ease of endoscopy, as both the time to reach the caecum, and pain perceived were highly concordant (proband concordance: 85% and 56% respectively). We found the expected positive relationship between BMI and colonoscopic anomalies such as diverticular disease and polyps in the whole population, but within twin pairs this association was reversed. This suggests that familial factors confound these associations. Host and microbial Next Generation Sequencing and metabolomics of the samples collected are planned in this cohort.

## Introduction

The colon, is the last part of the digestive system where water, salt and some vitamins, such as vitamin K or thiamine, are absorbed prior to defaecation. It is also a key location where microbial fermentation of remaining solid waste material takes place (1–3). The large intestine is populated by approximately 10^12^ microorganisms, out of the circa 10^14^ microorganisms hosted in different niches of the human body including skin, genitourinary and respiratory tracts, and small and large intestine (4–7). Over 700 different species live in the colon, prevalently dominated by Firmicutes and Bacteroides, with a varying ratio depending on different factors including health status (8–11). Interactions between the microbiota and the colon can be classified as mutualistic, symbiotic or pathobiontic (12, 13) and evidence is mounting for a role in host health and disease. Most human studies to date investigate the relationship between faecal samples and host physiology. However, animal studies have indicated tighter relationships between colonic microbiota and host physiology than with the stool, and highlighted the influence of microbiota on colonic gene expression (14).

ExHiBITT — Exploring Host microBlome in Teraction in Twins-is a sub-study within TwinsUK cohort (15, 16), which will enable scientist access to a large number of OMICS’ data related to the colon. Twin studies may particularly useful to study deep-tissue microbiota-host interactions, in part because of the strong influence of host genetics on gene expression and immune function disease associations, and to a lesser extent on microbiome itself (17–20). By analysing changes in monozygotic twins, with the same host genetics, effects of different microbiota can be examined without the variance attributable to host genetics. Thus, twins’ studies are recognised for their potential to investigate different phenotypes separating genetic from environmental effects (21). Monozygotic (MZ) twin pairs, where genetic variation is rare or null, provide the ideal scenario to investigate the effect of environmental factors such as gut microbiota, diet, smoking status or living habitat (22, 23). Analysis of samples, using high-throughput techniques, including Next Generation Sequencing (NGS), metabolomics and immune profiling of peripheral blood and caecum of twin pairs, is underway to investigate the host and microbiome genetics, metabolome and associated modulations of the immune system.

The objective of the present study was firstly to describe the distribution of this newly established cohort according to three different types of phenotypes: i) colonoscopy findings, ii) biomedical phenotypes and iii) environmental factors, and to assess the twin concordance for endoscopic variables. We then interrogated the relationships between BMI and colonoscopic findings using standard and within-pair regression modelling. Secondly, although routine endoscopic biopsies are considered safe, there is limited outcome data in patients that have large numbers of research biopsies taken, and where available is retrospective in nature (24). In this study we report on the safety and tolerability of taking more than 20 research biopsies within an older adult population.

Analysis of the colonoscopic findings found within this non-clinical population are not trivial. Polyps are tumours affecting approximately half of the western population at some point in life and detected in up to a third of all colonoscopies (25). The majority of polyps are adenomatous and, by definition, dysplastic with malignant potential. Adenomatous polyps increase with age, occurring in 21-28% 50-59 year olds, 41-45% in 60-69 year olds, and 53-58% in patients over 70 (26). Dysplastic polyps which are left undetected can develop into colorectal cancer (CRC), the third most prevalent cancer worldwide (27–30). There is interest, therefore, in understanding the development of polyps as a precursor of cancer. Diverticular disease is the symptomatic manifestation (normally abdominal pain) of people who have develop diverticula, which are small bulges in the large intestine (31). Approximately 1 every 4 people with will develop diverticulitis, which is the inflammation lead by bacteria and is associated with increased risk of intestinal perforation (32).

## Material and Methods

### Study ethical approval and participants consent

The ethics of this study were approved by the English National Health Service (NHS) Research Ethics Committee in June 2015. Participants provided informed written consent after registration and hold the right to drop out at any point of the study.

### Recruitment

The TwinsUK ExHiBITT — Exploring Host microBlome in Teraction in Twins-cohort was established between 2015 and 2018 to study interactions between colon microbiota and host genomics. Twins were recruited from the TwinsUK cohort with the eligibility criteria outlined in Supplementary_material_l.

Individuals who fell under this criterion were contacted by email. As the focus of our study was healthy ageing, individuals were recruited from older age bands preferentially.

### Data and sample collection

This cohort was annotated for three different types of phenotypes described in Supplementary_material_2.

Living area was assigned by extracting Land Cover Map (LCM) 2015 1 km target class for each of the participant’s postcode using R package ‘raster’ and ‘rgdal’. LCM classes were then reassigned as urban, suburban or rural. Phenotypes were assessed thought self-reported questionnaires in all cases except for weight and height in BMI, which were measured the day of the visit. SocioEconomic Status (SES) was based on postcode location and assigned using published deciles of the Index of Multiple Deprivation (IMD) for Scotland, Wales, England and northern Ireland, where 1 is the most deprived and 10 is the least deprived (33). Frailty index was annotated as described in Searle *et al*. (2008) (34).

Every patient underwent a colonoscopy, using the same bowel preparation (sennakot and sodium picosulphate). Colon biopsies were taken at colonoscopies from up to four locations (right colon, left colon, terminal ileum and cecum), caecal fluid, saliva and blood samples were collected at time of visit (Supplementary_material_3). Stool samples were taken 24 hours prior, before bowel preparation, and also at more than one week after the visit.

Data recorded just before commencing colonoscopy included presence/absence of irritable bowel syndrome (IBS), and presence/absence of a history of abdominal pain, loose stool or constipation. Phenotypic information collated during colonoscopy included endoscopic findings (i.e. polyps and location and number of areas containing diverticulae), pain scores as assessed by the endoscopist using the modified Gloucester scale (35) (1= comfortable, 5= frequent discomfort with significant distress), quality of bowel preparation and time to caecum. Histological outcomes from clinical biopsies of lesions were collated after the procedure.

### Immune profiling from peripheral blood and biopsies

Peripheral blood mononuclear cells (PBMC) were isolated using ficoll-paque density gradient centrifugation method. Multi-parametric flow cytometry was performed after staining with relevant fluorescent monoclonal antibodies to quantify T cell. Effector memory T-cells were identified as CD3^+^ CD4^+^ CD25^−^CD45RO^+^CD45RA^−^CCR7^−^, which then subsequently defined Th1 (CXCR3^+^CCR6^−^) Th17 (CXCR3^−^CCR6^+^), Th1-17 hybrid (CXCR3^+^CCR6^+^) and Th2 (CXCR3^−^ CCR6^−^CCR4^+^) cells. Antigen experienced regulatory T cells (Ag Exp Treg) were defined as CD3^+^CD4^+^ CD25^+^ CD45RA^−^CCR4^+^ which were then subdivided into T helper like subsets based on CCR6 and CXCR3 expression (Figure 1, panel a): 1A).

**Figure 1.**
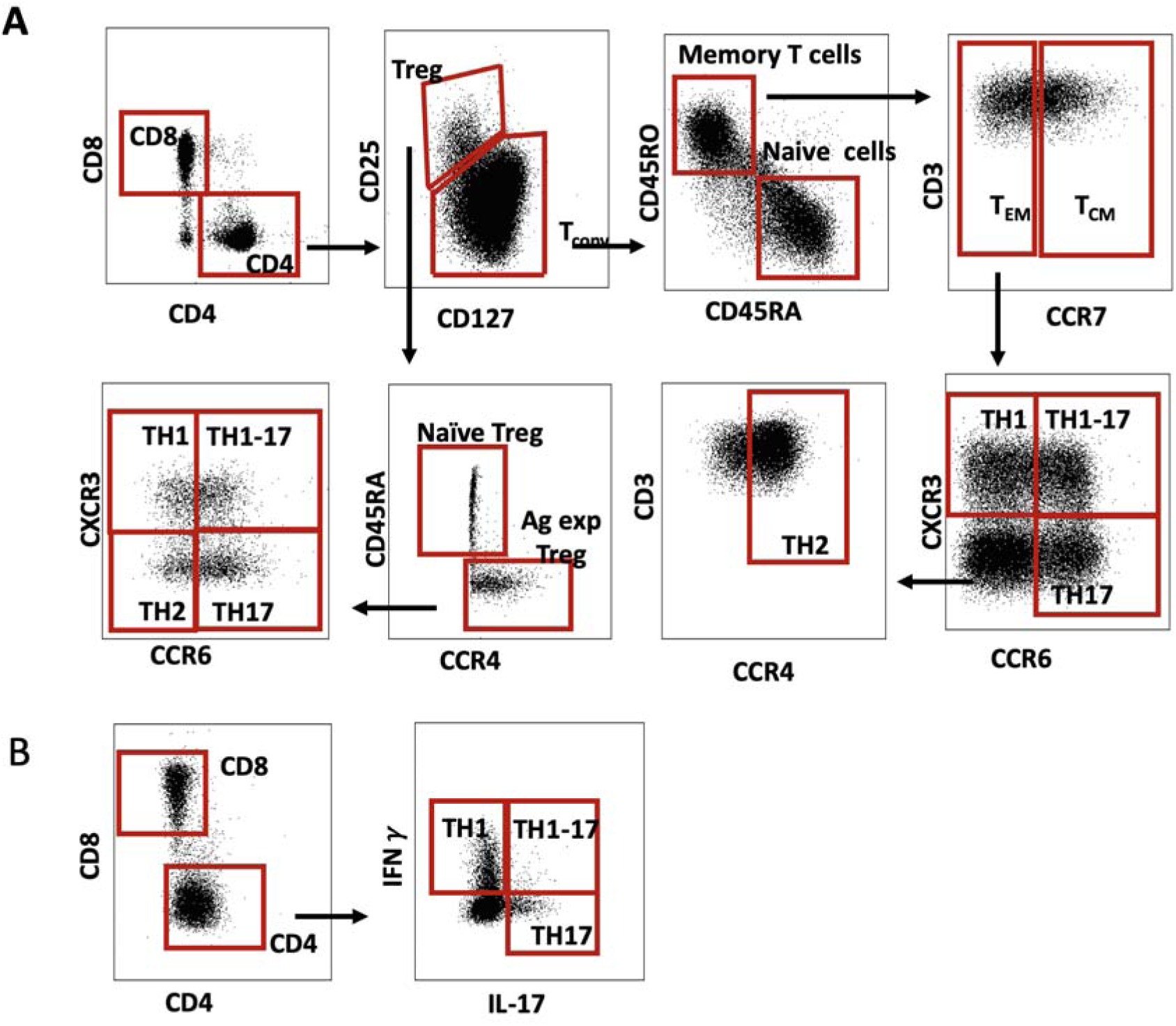
Flow cytometric gating strategy. Panel a) Flowcytometrie analysis of peripheral blood CD4 T cells (gated on CD3+ live lymphocytes) which were then divided into CD127^high^CD251^ow^ conventional T cells (Tconv) and CD127^low^CD25^high^ regulatory T cells (Treg). Tconv cells were then divided into naive and memory T cells. CD45RO^+^CD45RA^−^ memory T cells were subdivided into CCR7^−^ effector memory (TEM) and CCR7^+^ central memory (TCM) T cells. TEM defined Th17 (CCR6^+^CXCR3^−^), Th1 (CXCR3^+^CCR6^−^), Th1-17 (CXCR3^+^CCR6^+^) and Th2 (CXCR3^−^CCR6^−^CCR4^+^) cells. Antigen experienced Treg (Ag exp Treg) were defined as CD45RA^−^CCR4^+^ Treg which were then subdivided into T helper like subsets based on CCR6 and CXCR3 expression. Panel b) Flow cytometric analysis of lamina propria mononuclear cells-CD4 T cells (gated on CD45^+^CD3^+^ live lymphocytes) were divided into Th1, Th1-17 and Th17 cells based on IFN gamma and I1-17 expression.

Endoscopically acquired colonic biopsies were sampled and partially disrupted by gently compressing the epithelial/luminal aspect of the biopsy into the foam matrix. Complete culture medium (supplemented with rhIL2, broad spectrum antibiotics and anti-fungal reagents) was added and immune cells progressively migrated out of tissue into the culture medium. Cells were harvested after 48 hours for downstream analysis. Leukocyte yield using this system was typically in the region of 2×10^5^ cells per biopsy. The cells were then stimulated with PMY and ionomycin for 3 hours and analysed by intracellular cytokine staining and flow cytometry. T helper cell subsets were defined as Th1 (IFN-γ^+^IL-17^−^), Th17 (IFN-γIL-17^+^), and Th1-17 (IFN-γ^+^IL-17^+^) cells. (Figure 1, panel b).: 1B)

The data for each type of cell was calculated as a percentage of parent cell population and analysed using graphpad prism software.

### Statistical analysis

**Descriptive statistics** for sex, rearing, ethnicity, smoking status, living area and socioeconomical status as well as polyp presence) and measured variables (BMI, age and frailty) were calculated using RStudio (version 0.99.489 — © 2009-2015 RStudio, Inc).

For the concordance analysis of colonic traits, twin pairs where one of the individuals had missing information were removed. The formula employed was: *CR pairwise= (Number of concordant pairs* **/** *(Number of concordant pairs* **+** *number of discordant pairs)) *100* and *CR proband=(2*Number of concordant pairs / ((2*Number of concordant pairs)* **+** *number of discordant pairs*)) * 100.

**Inferential statistics** were employed to interrogate the cohort through Linear Mixed Effect Models (LMEM) using the algorithm provided in the R package lme4 (36). The model employed was: *lmer(Trait* **~** *Frailty* + *Age* + *BMI* + *Quality_of_bowel_prep* + *(1 | Family_No))*, where the random effects were the biological variates (frailty, age and BMI) and a technical covariate (quality of bowel preparation). The fixed effect was family relatedness. The traits studied were the four colonoscopy-derived phenotypes previously described. Moreover, a second model: *lmer*(*Time_to_caeum ~ Pain_score* + Endoscopist + AbdSym_including_lBS + Quality_of_bowel_prep + **Age** + Frailty + BMI + (1 | Family_No) was employed to identify any connexion between time to caecum and the phenotypes measured. Bonferroni correction was applied to all the results obtained from the statistical analysis.

Differences in *between* and *within* variation in twin pairs were studied using a linear model. The model used was: *lmer(Trait* ~ *BMI^b^+ BMI* + *Frailty^b^’* + *Frailty* + *(1 | Family_No))*, where ^*b*^ (between) denotes the mean for the trait in each family group, and ^*w*^ (within) the difference between individuals and the family mean for each pair. Statistical difference in the between and within coefficients for each trait was calculated using LINear COMbination of estimators (LINCOM), implemented in STATA, where the model was reiterated.

### Data availability

Data produced during the colonoscopy study will be publicly available through managed access. Researchers interested can request access following TwinsUK procedure available at TwinsUK Data Access Policy (http://twinsuk.ac.uk).

## Results and discussion

### 1. Recruitment

Two hundred and five twins volunteered for the study; out of those, two hundred successfully completed the colonoscopy. Withdrawals were linked to the discovery of a suspected cancer (n=3) or voluntary discontinuation during the intervention due to discomfort (n=2).

### 2. Samples collection

Colon biopsies were collected for interrogation of host genomics and microbiome analysis (data not reported here). Samples were conserved in liquid nitrogen and included biopsies from i) left colon (n=196), ii) terminal ileum (n=151), iii) caecum (n=73), and right colon (n=24) when one of the other locations was difficult to sample. Mucosal biopsies to be used for microbial analysis were conserved at −80°C. This included i) left colon (n=200), ii) right colon (n=179) and iii) caecum (n=79). Colon biopsies were taken in triplicates. In total, 5 replicates of caecal fluid were collected during the colonoscopy (n=197). Faecal samples were collected immediately prior to bowel preparation (n=169), and one week after (n=188). Other samples included saliva (n=180) and blood (n=204), which was stored as serum and plasma.

### 3. Cohort descriptive statistical analysis

The average age of the cohort was 58.70 ± 9.55 (F=58.60 ± 9.52, M=59.04± 9.38), BMI was 26.37 ± 5.22 (F=26.01±5.18, M=27.66±5.21), and frailty index 0.18±0.10 (F=0.19±0.10, M=0.17±0.09) (Supplementary_material_4, panel a). Twin pairs where differences between continuous traits (i.e. BMI, frailty index and EIMD deciles was bigger than 1 standard deviation were considered discordant (Supplementary_material_4, panel b). One twin pair was found discordant for BMI, and 2 for frailty. In total, 39 twin pairs were found discordant for EIMD decile. The twin pair with discordant BMI and frailty was also discordant for EIMD deciles.

In total, there were one hundred and sixty-one women and forty-four men in the cohort. Only four individuals (2%) were reared apart. Ninety-five percent of the individuals identified themselves as white, 2% as mixed, 2% as black and 1% as Asian. Five percent of twins could not attend the colonoscopy visit with their co-twins, and one individual’s twin dropped out from the study just before the visit. Smokers represented 26% of the cohort, 66% of individuals never smoked and 3% considered themselves as ex-smokers. Currently, 55% of the cohort live in the same county as their co-twin. Fifty-nine percent of the cohort live in sub-urban areas, 28% in rural areas and 11% in urban areas. Regarding socioeconomic status, 8% of the cohort were classified as belonging to IMD decile 1-2, 15% to SES 3-5, 42% to SES 6-8 and 35% to SES 9-10 where 10 is the least deprived (Supplementary_material_4, panel c).

In total, colonoscopy information from 196 individuals was collected. This information is next described following the time sequence of the data collection and from specific to accumulative phenotypic traits.

#### Pre-procedure outcomes

Twenty individuals (13%) reported irritable bowel syndrome (IBS), with a concordance rate of 50% (proband) respectively. Everybody with IBS reported one type or another of abdominal Symptom. The different types of symptoms recorded were: i) pain/cramps (n=26), ii) constipation (n=22), iii) rectal bleeding (n=2), diarrhoea (n=7) and alternative diarrhoea/constipation (n=6). The accumulative trait *‘presence of abdominal symptoms or IBS’* counted for 40 individuals (28%) affected by at least one symptom. The pairwise concordance rate was 48%, and the proband was 65% (Table 1). The influence of genetic and environmental factors on the emergence of IBS has been the subject of considerable debate, with increasing evidence that supports a role for genetic susceptibility. Our findings on concordance rates, is slightly higher than other MZ twin studies which have showed concordance rates between 17% and 33%(37–39).

**Table 1.**
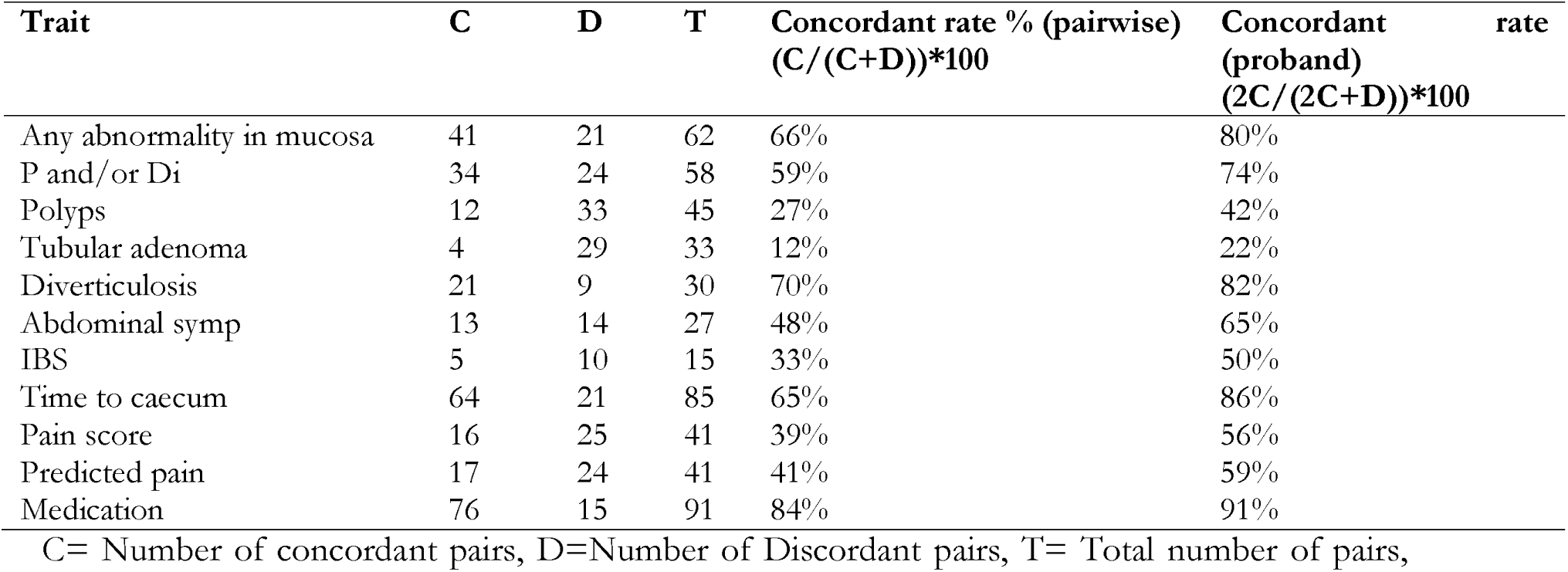
Concordance rate expressed in percentage.

#### Procedure related outcomes

Out of the 196 individuals with colonoscopy information, 4 of them had poor bowel preparation, 25 adequate and the rest had good bowel preparation. **Quality of bowel preparation** was used in the LMEM as a potential confounder.

**Sedation** provided included midazolam, fentanyl, endotox or nothing. The medication index was created considering the following factors: i) 1 mg of Midazolam=1 index unit, ii) 25 μcg of Fentanyl = 1 unit, and iii) Endotox use = 1 index unit. Sedation scores ranged from 1-6, average 3.8 ± 2.1. 15 twin pairs were discordant by more than one SD, giving a proband concordance rate equal to 91% (Table 1). These concordances should be taken with caution, as the endoscopist was not blinded to twin pairing. Concordant twin pairs for sedation were selected to study **pain scores** associations with time to caecum.

Two different types of pain traits were used, the original pain score taken during the colonoscopy and the predicted pain score adjusted taking into account the sedation, For that purpose, a Linear Mixed Effect Model (*Pain_score* **~** *Sedation_score* + *(1 |Family)*) was built in R to calculate the residuals from pain score taking into account family and sedation. Concordance rates were calculated in both traits, but only *Pain_score* of concordant twins for *Sedation_score* was used in the model to calculate associations with time to caecum. The minimum pain score was 0, and −1.19 in the predicted pain score. The maximum were 5 and 2.45 units respectively. The average pain score was 1.58 ± 0.79, and 0.003 ± 0.60 for the predicted pain score. The proband concordant rate for pain score was 56% and 59% for the predicted pain score (Table 1).

Time to caecum was in average 12.97 ±7.12 min, maximum time to caecum was 52 minutes and minimum 1.33. There were 64 concordant pairs and 21 discordant by more than one standard deviation. The concordance rate was 86% (proband)(Table 1). This minimal variation between twins in caecal intubation time, suggests that technical difficulty and by inference colonic morphology, was similar. Although this is not an entirely unexpected finding, it has not been previously described in MZ twin colonoscopies. Focussing only on those individuals with **polyps and/or diverticulosis**, in total 93 of them had one condition or both. Individuals had between 0 to 4 total polyps and/or diverticulosis in total (Supplementary_material_5, panels b, f and j). The proband concordance rate for polyps and/or diverticulosis was 74% (Table 1). This concordance is illustrated in Supplementary_material_6. Fifty-seven people had colonic **polyps** (28%), one of them had a potential cancer and appropriate actions were taken. The number of polyps ranged from multiple (>7) to 1 (average where present 1.5), (Supplementary_material_5, panels c, g and k). The concordance rate for polyps was 42% (proband). Only 4 pairs were concordant for tubular adenomas, the rest of the pairs discordant (n=29), giving a proband concordance rate of 22%.

Despite the fact that known **diverticular disease** was an exclusion for the study (due to the increased risk of bowel perforation), 51 people were found to have diverticulosis on endoscopy (26%), of which the majority (29) where located in the left colon. The number of locations for diverticulae within an individual ranged from 0 to 2. No individuals had evidence of inflamed diverticulae (diverticulitis). Twenty-one twin pairs (n=42) were concordant for diverticulosis and nine cases (n=9) were discordant (Fig 3, panels: I, j, k, and i). Thus, *diverticulosis* had the highest concordance between twins at 82% (proband) (Table 1, Supplementary_material_6).

In a previous twin cohort study from the Swedish Twin Registry, the MZ concordant rate for diverticulosis was 6% only, due to the fact there were over fourteen fold times more discordant twins for diverticulosis than concordant ones (40). Similarly, the diverticulosis study from the Danish Twin Registry found that the diverticulosis twin concordance rate was 8% (40, 41). Differences between these studies and the results from the colonoscopy TwinsUK is most likely to be a function of ascertainment. Our participants were selected not to have a known diagnosis of diverticulosis, and presence was ascertained endoscopically. Whereas these other studies relied on health record data from physician diagnosis and asymptomatic co-twins may not have undergone a colonoscopy. Alternatively there could come from environmental and genetic variation between Scandinavian and British populations, or differences in advances in the colonoscopy techniques (where employed), cohort size and recruitment criteria and timing of the study (the Swedish Twin Registry took data from 1886 to 1980, and the Danish went from 1977 to 2011, while the TwinsUK colonoscopy study examined volunteers between 2015 and 2018). Heritability of diverticular disease has been estimated by Strate and colleagues (2013) (42) as 53%, which could be an underestimate due to asymptomatic disease. To the best of our knowledge, the high endoscopic concordance rate for diverticulosis in identical twins identified in this cohort was never reported before. This indicates that genetic variants could contribute to the development of diverticulosis, as previously indicated.

##### Complications

Despite the large number of samples collected, there were no major complications, including perforation or bleeding. Minor complications included incomplete procedures secondary to patient discomfort (n=2) or presence of a fixed sigmoid that limited endoscopic progression. One 61year-old patient who received sedation experienced a transient vasovagal episode during the procedure.

#### Post-procedure related outcomes

One hundred and four individuals had some sort of **abnormality** in the mucosa observed either during the colonoscopy or at histology. Individuals with presence of any abnormality represented 51% of the cohort, 45% of the cohort was absent of any sort of abnormality and the remaining are those individuals with no colonoscopy information available (N/A). Abnormalities ranged from 0 to 4 (Supplementary_material_5, panels a, e and i). Twenty-one twin pairs (n=42) were discordant for any abnormality, and 41 twin pairs were concordant. This gave a concordant rate of 80% (proband).

### 4. Cohort inferential statistical analysis

A LMEM was used to interrogate if our colonoscopy traits were associated with biological covariates (i.e. BMI, age, frailty). Results from the LMEM showed that age and BMI were statistically significant according to i) *total number of abnormalities*, ii) *total number of polyps and diverticulosis*, and iii) *total number of polyps.* **To**tal number of diverticulosis was only relatively closed to be significant in the case of BMI (Supplementary_material_7, Table 2). There was no detectable association between time to caecum and biological covariates.

**Table 2:** Results from the LMEM used to interrogate the phenotypes from the colonoscopy analysis according to the biological covariates: BMI, age and frailty:

| Number of Abnormalities (n=190) | Estimate | Std. | t value | Pr(>Chisq) |
| --- | --- | --- | --- | --- |

|  |  | Error |  |  |
| --- | --- | --- | --- | --- |
| Frailty Index | -1.09 | 0.68 | -1.61 | 0.10 |
| Age | 0.03 | 0.01 | 3.60 | *** |
| BMI | 0.04 | 0.01 | 3.15 | ** |
| **Number of Polyps and Diverticulosis (n=190)** |  |  |  |  |
| Frailty Index | -0.97 | 0.61 | -1.60 | 0.11 |
| Age | 0.03 | 0.01 | 4.04 | *** |
| BMI | 0.05 | 0.01 | 4.04 | *** |
| **Number of Polyps (n=188)** |  |  |  |  |
| Frailty I | -0.81 | 0.52 | -1.56 | 1 |
| Age | 0.02 | 0.01 | 3.30 | ** |
| BMI | 0.03 | 0.01 | 3.16 | ** |
| **Number of Diverticulosis locations (n=183)** |  |  |  |  |
| Frailty I | -0.37 | 0.28 | -1.33 | 0.19 |
| Age | 0.01 | 0.01 | 2.02 | * |
| BMI | 0.01 | 0.01 | 1.52 | 0.13 |

Furthermore, between family (*b*) and within pair (*w*) twin differences for BMI and frailty^7^ index were studied using linear models. No significance was found for frailty. Reflecting the results above, and consistent with previous published studies (43–46), BMI*b* was statistically significant in all the traits studied such that higher BMI led to greater risk of anomalies. BMI difference within pairs was significantly different from the between family difference in all four tests and showed significant opposite relationship in the traits i) *total number of polyps*, and ii) *total number of polys and/or diverticulosis*, i.e. higher BMI within pairs led to reduced risk of anomalies. This could indicate common factors to both twins, such as genetics and early life environment, could be the link between with BMI and the colonoscopy traits studied such as polyps (Table 3), rather than BMI being directly causal. This is intriguing given the evidence of host genetic factors impacting the gut microbiome (47), and obesity^7^ (48). Only a minority^7^ of studies have looked at microbiome as a potential biomarker associated with the development of polyps in healthy individuals (30, 46, 49). Further work with ExHiBITT will consider microbiome composition in relationship to polyps and diverticular disease.

**Table 3:**
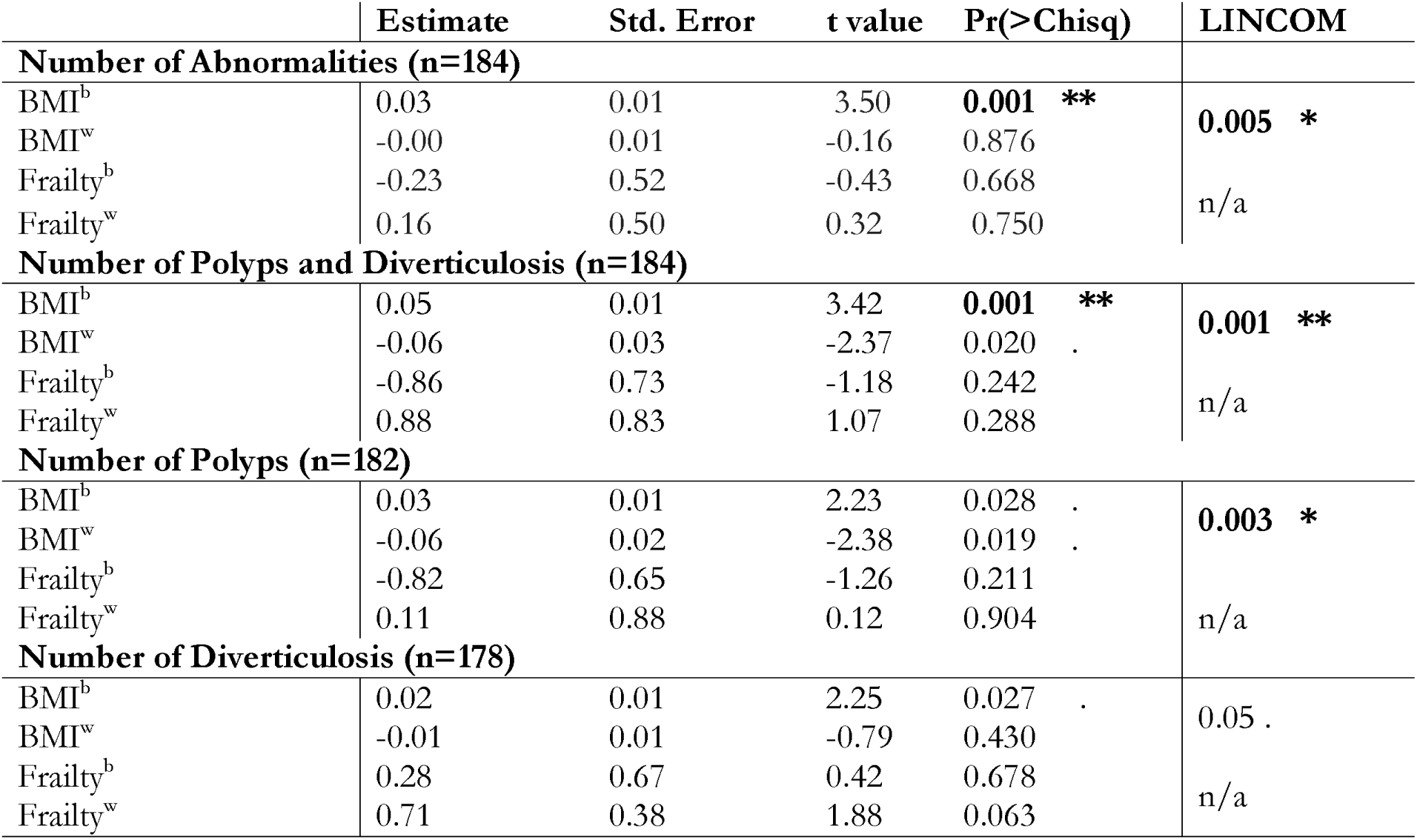
Results from the LMEM and LINCOM test used to interrogate the between and within variation in BMI and frailty:

#### Immune profiling outcomes

The twin pairs were highly concordant for different T cell subsets in both blood (Figure 2, panel a) and gut (Figure 3, panel a). Preliminary analysis showed differences in the immune response between males and females such as increased CD4 proportion and reduced antigen experienced Treg in females (not shown). Interestingly we found increased proportion of Th17 and Th2 cells in the peripheral blood in autumn-winter seasons compared to spring-summer seasons (Figure 2, panel b). No marked differences were seen in the peripheral blood immune profile in traits such as polyp or diverticulosis. However, increased proportion of hybrid Th1-17 cells producing both IFN gamma and IL-17 were found in colonic biopsies from patients with diverticulosis (Figure 3, panel b). No differences were found in the gut immune profile of individuals with/without polyps. Since generation of effector T-cell responses has been shown to be dependent on the composition of the intestinal microbiota, it will be interesting to look at the microbiome driving these differences in our cohort.

**Figure 2.**
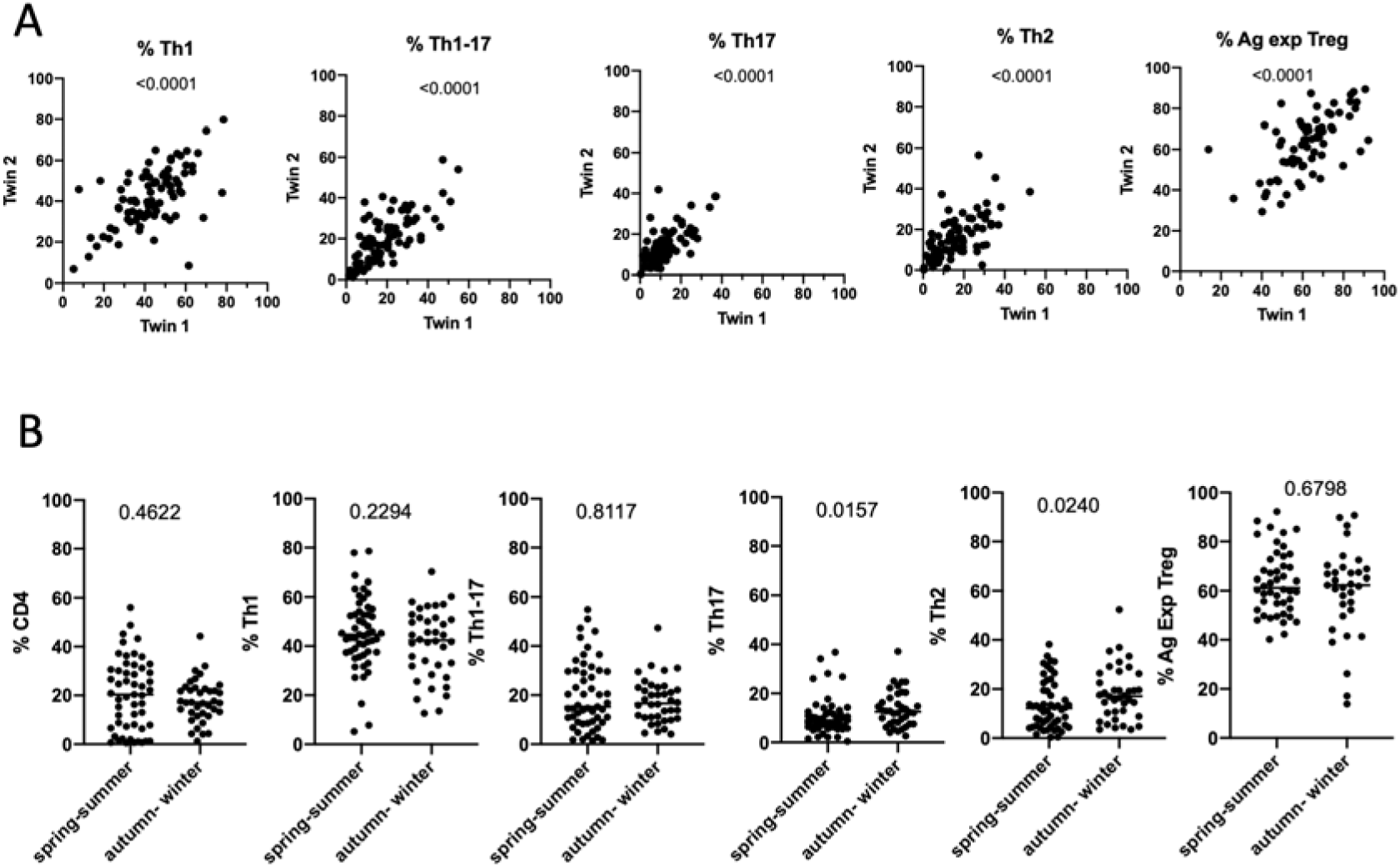
Peripheral blood immunophenotyping. Panel a) Proportion of different T helper cell subsets correlate between individual Twin pairs. Panel b) Frequency of CD4 T cells, Th1, Th1-17, Th17, Th2 and Ag exp Tregs between samples collected at different seasons.

**Figure 3.**
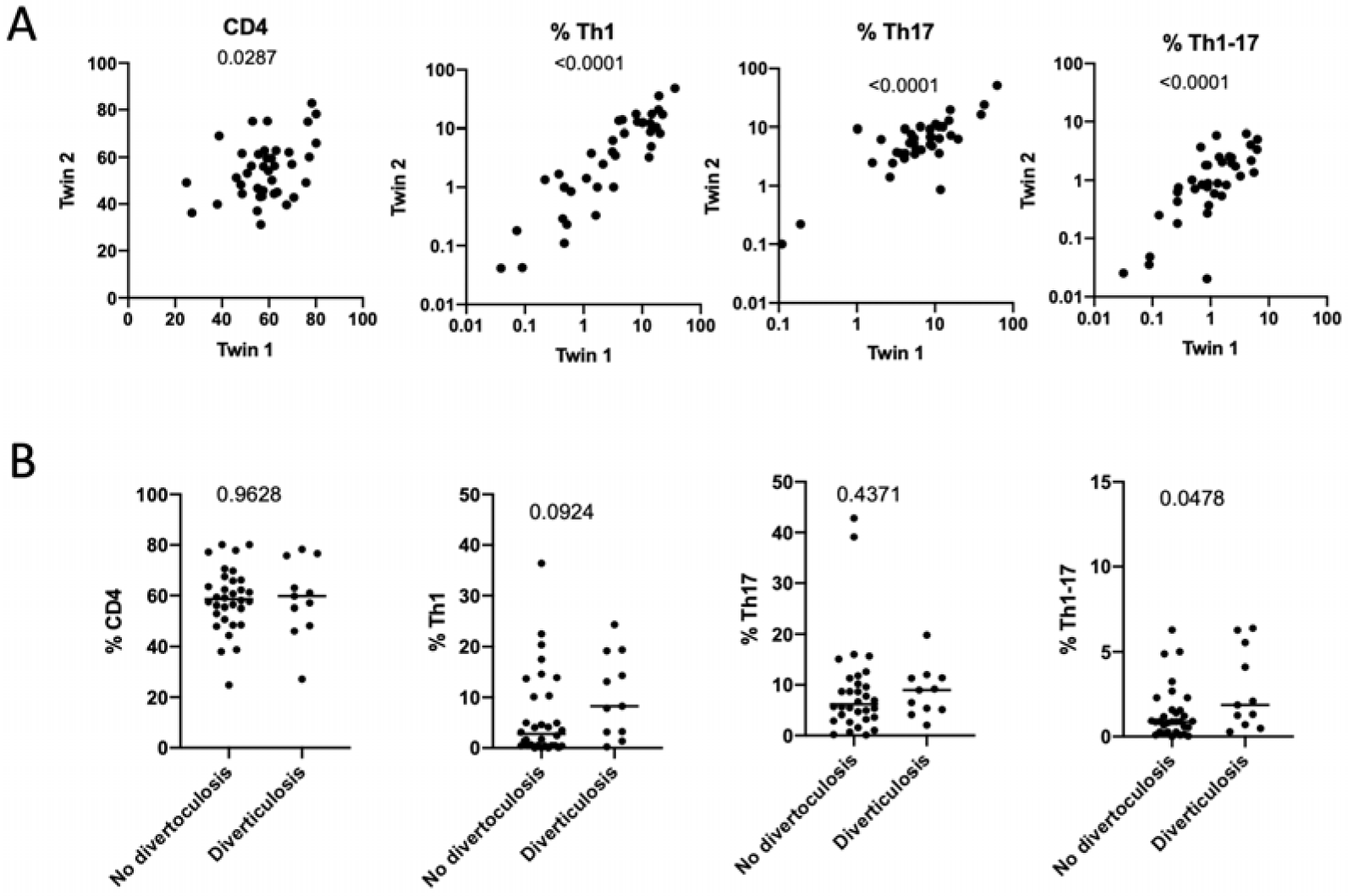
Gut immunophenotyping. Panel a) Proportion of different T helper cell subsets in the gut correlate between Twin pairs. Panel b) Proportion of CD4 T cells, Th1, Th1-17 and Th17between individuals with or without diverticulosis.

## Conclusions

This cohort represents a great potential to study microbiome-host interactions in the colon, and their implications for the host immune system. The cohort is annotated for a large number of phenotypes representative of UK society. Preliminary findings showed that polyps are strongly correlated with BMI and age, but that the relationship with BMI may be confounded by factors genetics and other factors shared by twins. There is a high rate of concordance between twin pairs for diverticulosis, less so for polyps. Interestingly, similar intubation times and pain scores were found for twin pairs, which could indicate that familial factors determine the ease of colonoscopy for both the endoscopist and patient. Further studies will include the high throughput analysis of the samples. We have also successfully phenotyped immune profile from the blood and gut of healthy twin pairs. High rate of concordance was found among twin pairs for effector and regulatory T cell subsets highlighting genetic control of immune response in monozygotic twins whereas seasonal variations found in the proportion of effector cell subsets ascertains the environmental programming of immune responses. Hybrid Th1-17 cells in the gut were shown to be associated with diverticulosis. Further analysis of this cohort will reveal the ileal microbiota responsible for driving systemic and mucosal immune response.

## Abbreviations

BMI: Body Mass Index
MZ: Monozygotic
NGS: Next Generation Techniques
NHS: National Health Service
SES: socioeconomic status

## Declarations

The authors declare they do not have conflict of interest.

## Acknowledgments

The authors thank the twins for their participation in the study, and the Medical Research Council (MRC) for funding this research (RE10740). We also wish to thank Clare Stockwell, Rachel Horsfall and Isabelle Granville Smith from the Microbiome Project, and Genevieve Lachance, Darioush Yarand and Merve Demirol from IT/Data & Administration resources, King’s College, University of London, for their technical assistance. Finally, we would also like to thank to Dr Julia El-Sayed Moustafa for the advice provided with the statistical models.

## Authors’ contributions

MMO wrote the first draft of the manuscript, compiled the metadata and created the figures. HI contributed to collect colonoscopy data and contribute to the gastroenterological aspects of the manuscript. SHK carried out all the immunological analysis and contribute to write the manuscript. RB compiled the metadata related to socioeconomic status, frailty and geographical location. NP conducted the colonoscopies. TS, KS and CS conceived the idea and supervised the work. All authors contributed to the experimental plan, supervised the work and contributed to write the manuscript. All authors have approved the final manuscript.

