## Supplementary_material_1 for "Introducing ExHiBITT – Exploring Host microbiome inTeractions in Twins-, a colon multiomic cohort study"

Box 1.

| **Box 1- Eligibility criteria for** **TwinsUK ExHiBITT** | |
| --- | --- |
| **Inclusion** | - Identical (MZ) twins - Both twins willing to participate in study |
| **Exclusion** | - History of type 1 diabetes, rheumatoid arthritis, inflammatory bowel disease, colon cancer or diverticular disease - On anti-platelet or anti-coagulant medications that cannot be safely stopped |
