## Supplementary_material_2 for "Introducing ExHiBITT – Exploring Host microbiome inTeractions in Twins-, a colon multiomic cohort study"

Box 2.

| **Box 2- Phenotypes annotated in TwinsUK ExHiBITT** | |
| --- | --- |
| **Biomedical phenotypes** | - Sex - Ethnicity - Rearing - Age - BMI - Frailty |
| **Environmental factors** | - Smoking status (smokers, ex-smokers and non-smokers) - Geography (rural/suburban/urban, concordance by county) - Socioeconomical status - Visit details (paired or separate) |
| **Colonoscopy findings** | - Polyps - Diverticulosis - Normal mucosa - Quality of bowel preparation for the colonoscopy - Time to caecum and endoscopist - Abdominal symptoms (i.e. pain, constipation, rectal bleeding, diarrhoea) and presence of IBS. - Sedation used and pain scores. |
