## Supplementary_material_3 for "Introducing ExHiBITT – Exploring Host microbiome inTeractions in Twins-, a colon multiomic cohort study"

**Supplementary material figure 1**

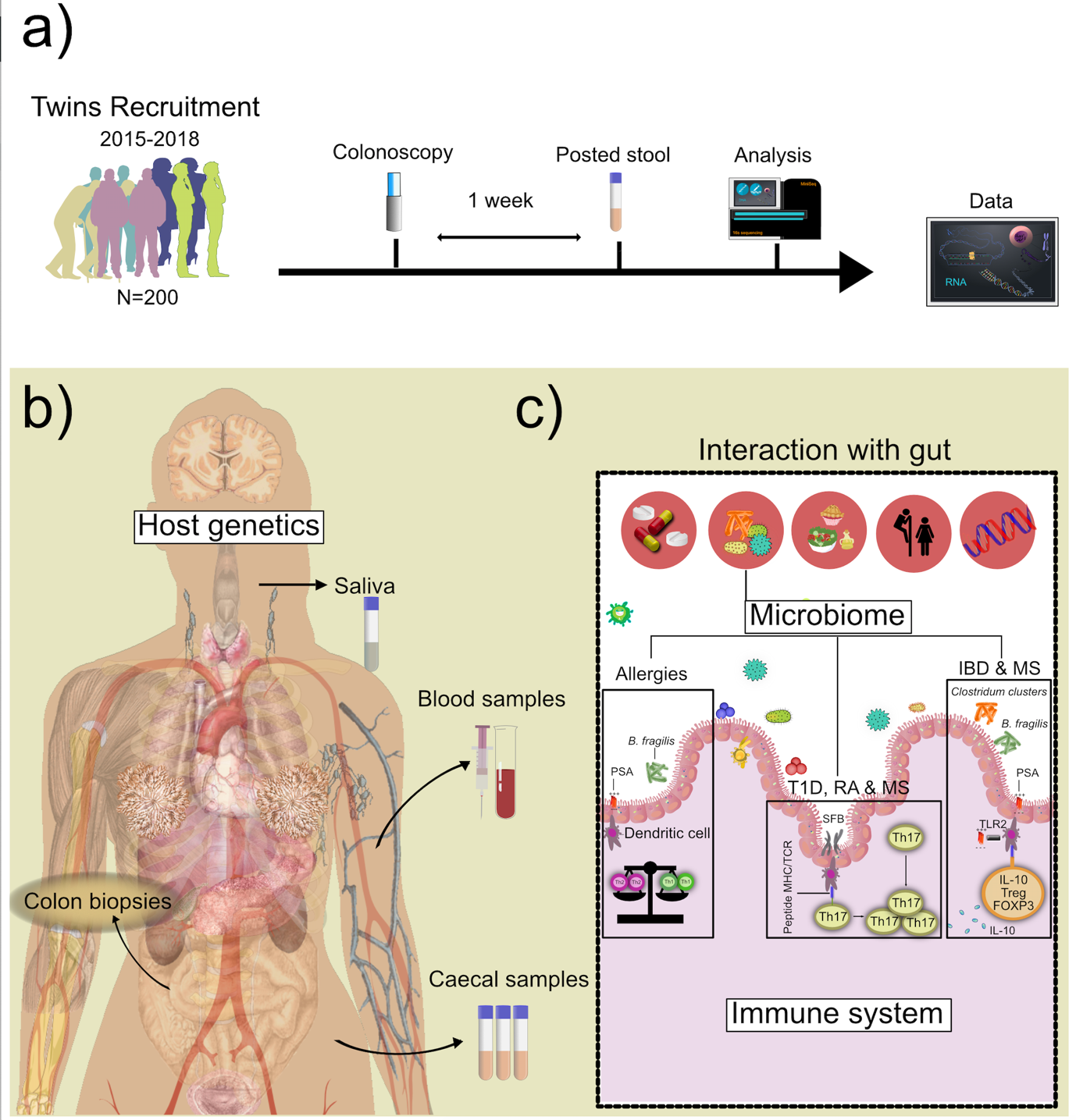

**Supplementary Material Figure 1.** Panel a) Study design included the recruitment of at least 200 twin volunteers. All samples were taken on the day of the visit except for the stool posted sample that was donated one week after. Panel b) Samples taken on the visit included colon biopsies, blood, saliva and caecal fluid. Panel c) Different factors can interact with the host immune system, among them the microbiota. Some of the mechanisms previously described involved allergies, Type 1 Diabetes (T1D), Rheumatoid Arthritis (RA), Multiple Sclerosis (MS) and Inflammatory Bowel Disease (IBS). *Panel c) was adapted from (12)*.
