## Supplementary_material_4 for "Introducing ExHiBITT – Exploring Host microbiome inTeractions in Twins-, a colon multiomic cohort study"

**Supplementary material Figure 2.**

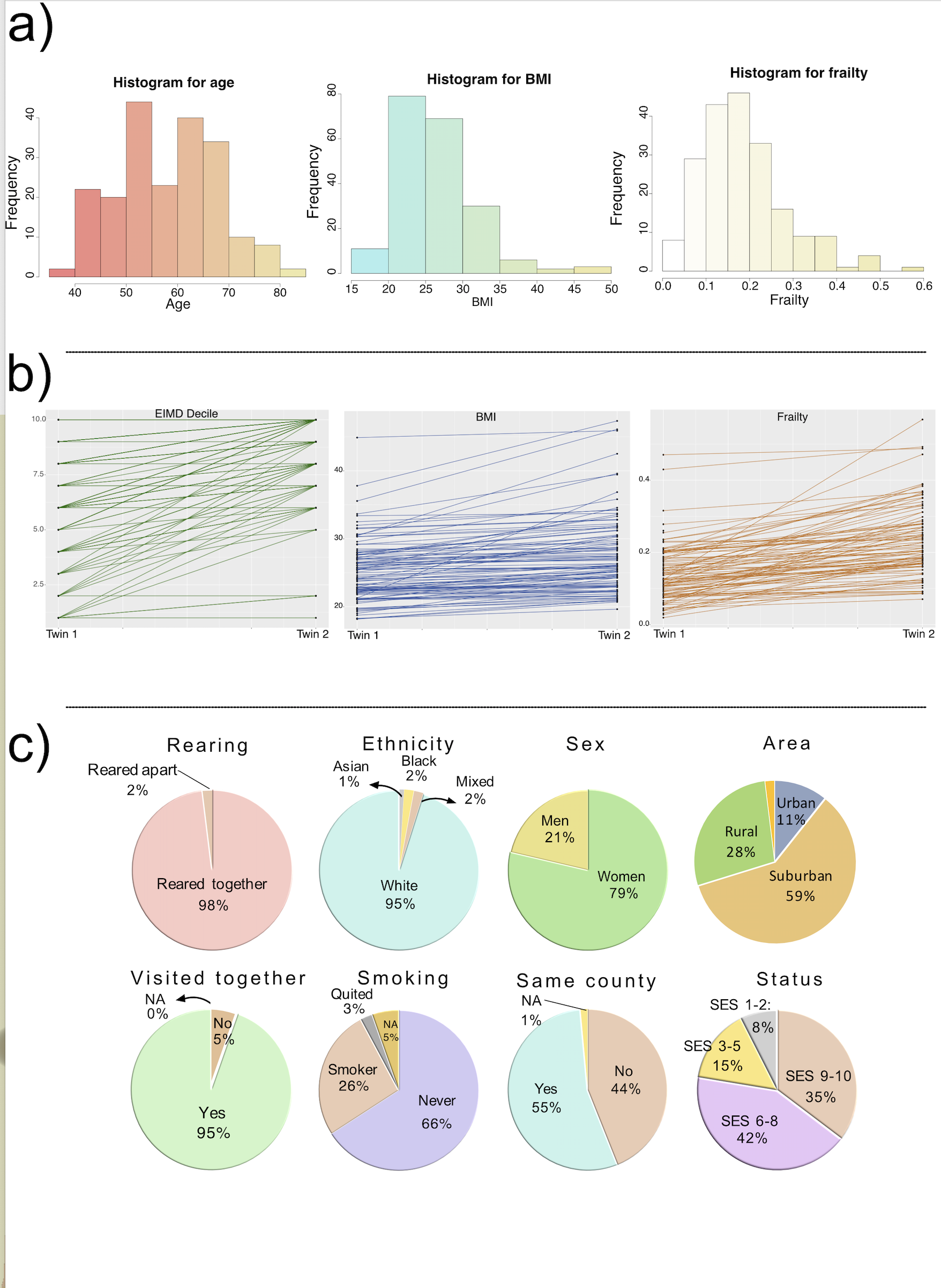

**Supplementary material Figure 2.** Cohort distribution. Panel a) Histograms with population distribution for age, BMI and frailty (from left to right). Panel b) Line chart pairing twin one (lowest value for each trait) with twin two (highest value for the trait). Steep lines indicate discordancy between traits. These traits are, from left to right: socioeconomical status, BMI and frailty. Panel c) Pie charts showing percentages of individuals belonging to each class of the phenotypes: rearing, ethnicity, sex, area where are they living, whether or not the visited together the colonoscopy appointment, smoking status, whether or not they are living in the same county and socio-economic status (SES).
