## Supplementary_material_5 for "Introducing ExHiBITT – Exploring Host microbiome inTeractions in Twins-, a colon multiomic cohort study"

**Supplementary material figure 3.**

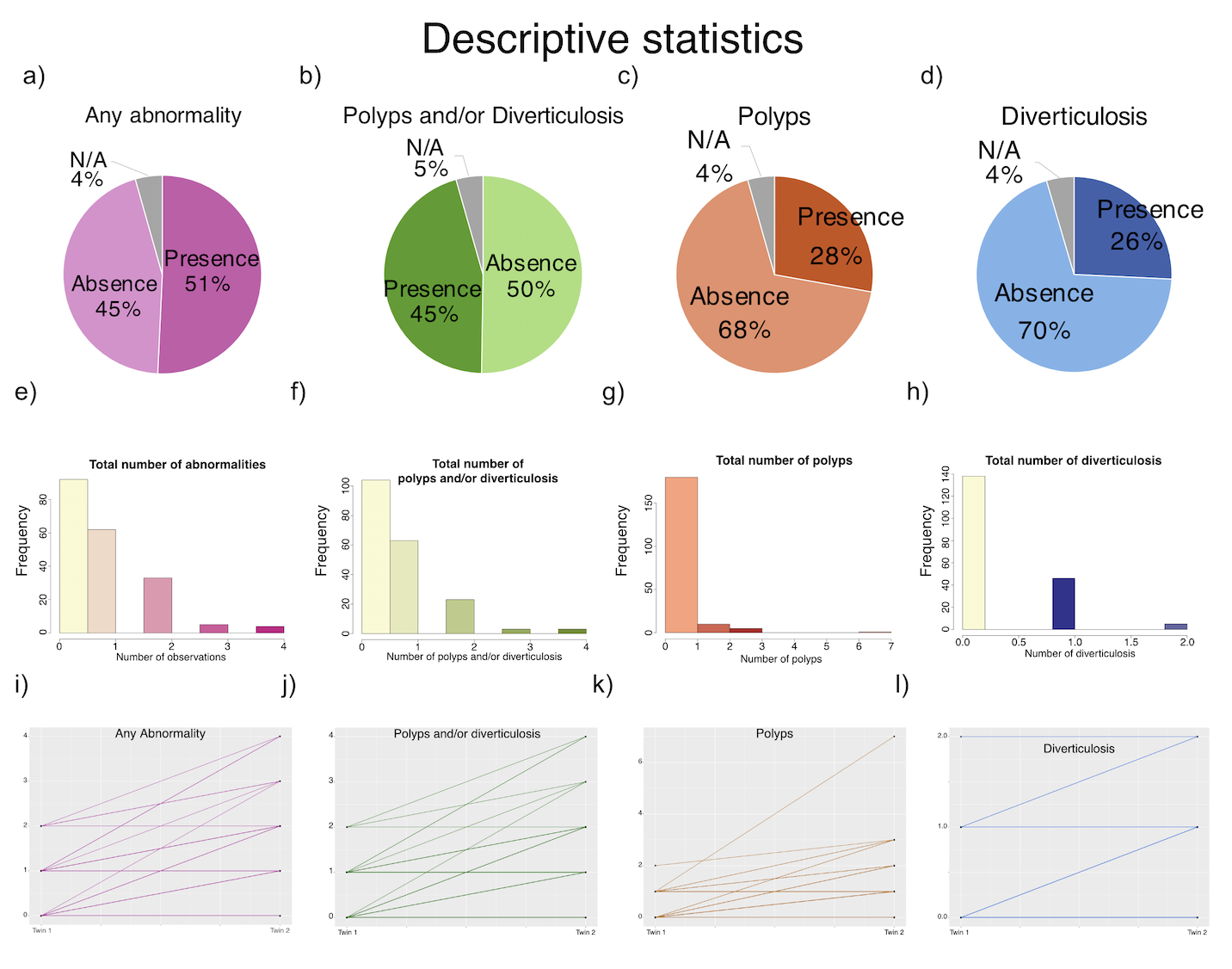

**Supplementary material figure 3.** Colonoscopy descriptive statistic. In columns from left to right the phenotypes are: i) number of abnormalities in mucosa, ii) number of polyps and/or diverticulosis locations, iii) number of polyps and iv) diverticulosis locations. Panels: a, b, c, and d: pie-charts representing the percentages of presence/absence of the phenotypes. Panels: e, f, g and h: histograms representing the distribution of the cohort. Panels i, j, k, and l: line charts representing discordant phenotypes.
