## Supplementary_material_6 for "Introducing ExHiBITT – Exploring Host microbiome inTeractions in Twins-, a colon multiomic cohort study"

**Supplementary material figure 6.**

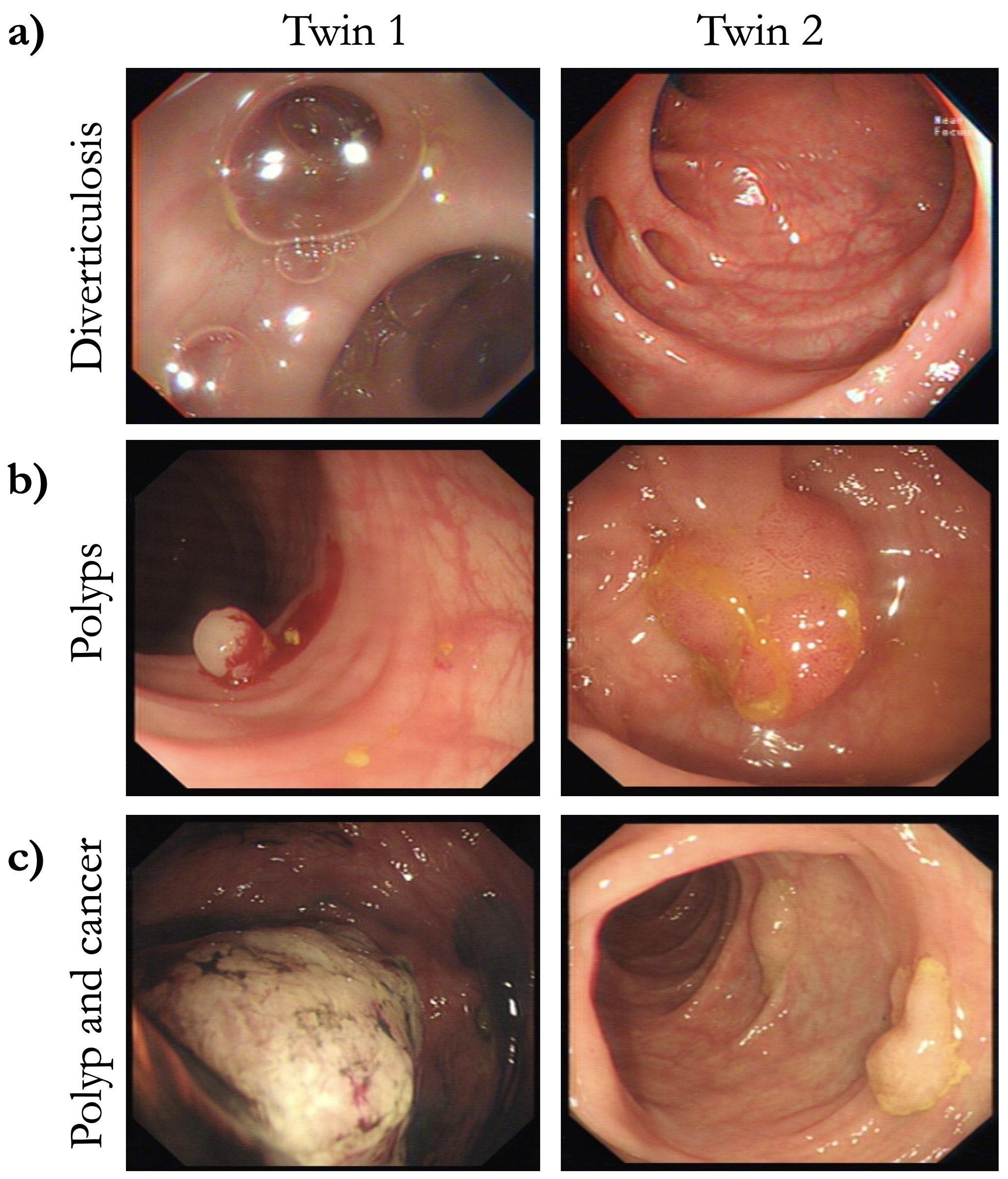

**Supplementary material figure 6.** Concordant twin pairs for: a) diverticulosis and b) polyps. c) Twin pair where one twin had a potential cancer and twin 2 had more had 7 polyps.
