## Supplementary_material_7 for "Introducing ExHiBITT – Exploring Host microbiome inTeractions in Twins-, a colon multiomic cohort study"

**Supplementary material figure 5**
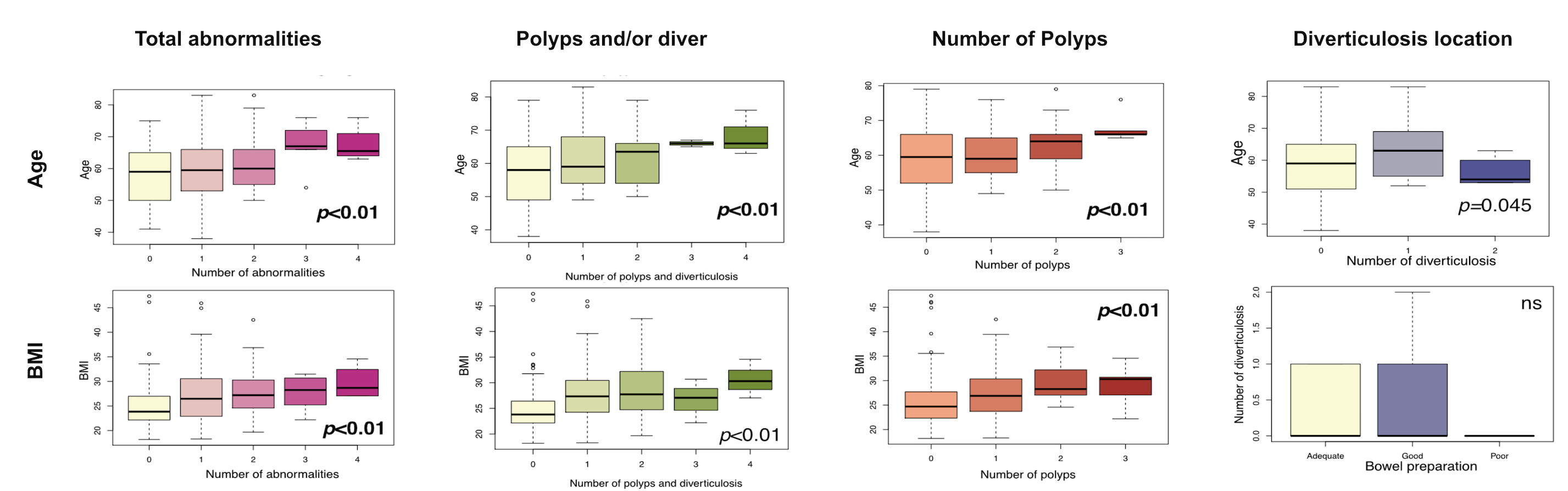
 **Supplementary material figure 5.** Colonoscopy inferential statistics. Boxplots in panels from left to right show the traits: total number of abnormalities in mucosa, number of polyps and /or diverticulosis location, total number of polyps and total number of diverticulosis locations. The top row shows effects in age, and the bottom one in BMI. Frailty was not found significant.
